# Evolutionarily Conserved Alternative Splicing Across Monocots

**DOI:** 10.1101/120469

**Authors:** Wenbin Mei, Lucas Boatwright, Guanqiao Feng, James C. Schnable, W. Brad Barbazuk

## Abstract

One difficulty when identifying and analyzing alternative splicing (AS) events in plants is distinguishing functional AS from splicing noise. One way to add confidence to the validity of a splice isoform is to observe that it is conserved across evolutionarily related species. We use a high throughput method to identify junction based conserved AS events from RNA-Seq data across nine plant species including: five grass monocots (maize, sorghum, rice, *Brachpodium* and foxtail millet), plus two non-grass monocots (bananan and African oil palm), the eudicot *Arabidopsis* and the basal angiosperm *Amborella*. In total, 9,804 conserved AS events within 19,235 genes were identified conserved between 2 or more species studied. In grasses containing large regions of conserved synteny, the frequency of conserved AS events is twice that observed for genes outside of conserved synteny blocks. In plant-specific RS and RS2Z subfamilies, we observe both conservation and divergence of AS events after the whole genome duplication in maize. In addition, plant-specific RS and RS2Z subfamilies are highly connected with R2R3-MYB in splicing networks. Furthermore, we discovered that the network based on genes harboring conserved AS events is enriched for phosphatases, kinases and ubiquitylation genes, which suggests that AS may participate in regulating signaling pathways. These data lay the foundation for identifying and studying conserved AS events in the monocots, particularly across grass species, and this conserved AS resource identifies an additional layer between genotype to phenotype that may impact future crop improvement efforts.

## Introduction

Alternative splicing (AS) is an important post-transcriptional process that can produce two or more transcript isoforms from a pre-mRNA. AS occurs in the spliceosome by removing introns and joining exons through the selective use of splice sites (Lee and Rio 2015) and is governed by *cis-* regulatory elements such as splicing enhancers/silencers, and *trans-* regulatory elements including SR proteins and hnRNP proteins (Busch and Hertel 2012, Wang and Burge 2008, Reddy et al. 2012, Reddy et al. 2013). AS isoforms that are translated can influence proteome diversity, while others that are not translated are purposefully non-functional and can act to post-transcriptionally modulate gene product levels (Fu et al. 2009, Hammond, Wachter, and Breaker 2009). AS participates in many important processes during the lifecycle of plants (Staiger and Brown 2013) and AS occurs in response to many abiotic stresses (Mastrangelo et al. 2012), for example red/blue light (Shikata et al. 2014, Wu et al. 2014), salt stress (Feng et al. 2015), drought (Thatcher et al. 2016), flooding (Syed et al. 2015) and temperature (Streitner et al. 2013, James et al. 2012).

RNA-Seq has become a standard tool to investigate transcriptomes and AS isoforms. While many studies have identified AS in individual species (Li, Xiao, and Zhu 2014, Thatcher et al. 2014, Shen et al. 2014, Filichkin et al. 2010, Mandadi and Scholthof 2015), few report investigating conserved AS across multiple plant genomes. Most published studies have focused on identifying conserved AS events between only two species. For example Severing et al. (2009) identified 56 protein-coding, conserved AS events between *Arabidopsis* and rice orthologous gene sets, and 30 conserved AS events leading to non-functional isoforms subjected to nonsense-mediated decay. Similarly, another study detected 537 AS events conserved between *Arabidopsis* and *Brassica* (Darracq and Adams 2013) and 71 conserved AS events were identified between *Populus* and *Eucalyptus* (Xu et al. 2014). With respect to comparing more than two species, one study found 16 AS transcripts conserved between *Brachypodium*, rice, and *Arabidopsis* by performing all-vs-all BLAST between EST sequences (Walters et al. 2013). Recently, discovery and identification of AS events conserved broadly across eudicots has been reported (Chamala et al. 2015). Chuang et al. (2015) used available Sanger EST sets to characterize AS in grass species but did not focus on conserved AS events. There are no corresponding multi-species studies that leverage the deep NGS resources to characterize and identify conserved AS events within monocots and the grass family.

Many monocot species are economically important and grass species (Poaceae) are a particularly important source of calories. Within the grass family there are two major clades. The BEP clade is made up of the Bambusoideae, Ehrhartoideae, and Pooideae and contains rice and *Brachypodium* (Zhao et al. 2013); the PACMAD clade includes Panicoideae, Arundinoideae, Chloridoideae, Micrairoideae, Aristidoideae, and Danthonioideae, and includes maize, sorghum and foxtail millet (Cotton et al. 2015). In addition to these two major clades, there are three clades of basal grasses lineage Anomochlooideae, Pharoideae, and Puelioideae (Cotton et al. 2015). Published whole genome sequences exist for many monocot species including the grass species maize (Schnable et al. 2009), sorghum (Paterson et al. 2009), rice (Goff et al. 2002), *Brachypodium* (Vogel et al. 2010) and foxtail millet (Bennetzen et al. 2012), the non-grass species banana (D’Hont et al. 2012) and African oil palm (Singh et al. 2013).

To characterize AS and identify conserved events due to species shared ancestral a computational method was employed that identifies high confidence isoforms in any species with RNA-Seq data and a reference genome sequence, and then uses a junction based approach to identify conserved AS events across multiple species. In this analysis, we focused on seven monocot species that have available reference genome sequences and deep RNA-Seq data sets. Five of these are grasses (maize, rice, sorghum, foxtail millet and *Brachypodium*) that also have substantial retained gene collinearity which eases the identification of synteny and aids orthologue calls (Schnable, Freeling, and Lyons 2012). This analysis also includes banana and African palm oil, which extends the discovery of conserved events into non-grass monocot species. Although there are other non-grass monocot species that have available reference genomes, such as duckweed and orchid, there was not sufficient RNA-Seq data available for these species to be included in this analysis. Likewise, species representing basal grass lineages were excluded due to lack of reference genome sequences. Finally, the eudicot *Arabidopsis thaliana* and the basal angiosperm *Amborella* were included to provide outgroups for comparison. Therefore, the species studied allowed us to identify conserved AS events at specific reference points across the grasses and extends into non-grass monocots, eudicots and the sister species to all angiosperms.

Examination of conserved events uncovers likely functionally important AS isoforms of SR protein genes, stress-response transcription factors, and members of known protein-protein interaction networks. In addition the conserved AS event collection provides an opportunity to test the functional sharing hypothesis (Su et al. 2006) between single-copy vs. non-single-copy genes, something that was not examined in previous studies of conserved AS events. This hypothesis predicts more instances of conserved alternative splicing events will be found in single copy genes than in non-single-copy genes. To measure to extent of selection pressure on AS, we examined Ka/Ks values on exons that flank conserved AS events. We also present network analysis based on the *Arabidopsis* protein-protein interaction data that provides evidence for a close association between splicing factors and stress responsive transcription factors that undergo conserved AS, and between splicing factors and R2R3-MYB across species. Finally, we discuss several noteworthy cases of conserved AS events across angiosperms including SPA3 possessing a conserved alternative donor site, a KH RNA-binding domain containing protein with a widely conserved exon skipping event, and the stress response NAC transcription factor RD26 that harbors an intron retention event conserved across the angiosperms.

## Materials and Methods

### Genome Assemblies and Annotations

All analyses were performed on publicly available sequence and annotation collections for the following species: foxtail millet (*Setaria italica*), rice (*Oryza sativa L. ssp*. *Japonica*), *Brachypodium* (*Brachypodium distachyon*), the non-grass monocots banana (*Musa acuminata*) and African oil palm (*Elaeis guineensis*), the eudicot *Arabidopsis thaliana* and the basal angiosperm *Amborella* (*Amborella trichopoda*), which is a sister taxon to all angiosperms.

The Maize RefGen_v2 genome sequence and working gene set annotation were retrieved from MaizeGDB (Lawrence et al. 2004). Reference genome sequence and gene annotation for oil palm were retrieved from the Malaysian Palm Oil Board (genomsawit.mpob.gov.my). Reference genome and annotation for Banana were retrieved from the banana genome hub (Droc et al. 2013). All remaining reference genome assemblies and annotations were retrieved from Phytozome v10 (Goodstein et al. 2012) (Table S1).

### Transcriptome Data Collection

AS in maize (*Zea mays*) and sorghum (*Sorghum bicolor*) was previously identified based on public RNA-Seq data and described in (Mei et al. In Review), and available from figshare (http://10.6084/m9.figshare.4205079). Transcript sequences and annotations of *Arabidopsis thaliana* were retrieved from Araport11 PreRelease 20151202 (Cheng et al. 2016), which presents the updated TAIR10 annotation by inclusion of 113 RNA-Seq data sets. For the remaining 6 species, Illumina RNA-Seq, 454 and Sanger EST transcript sequences were downloaded from NCBI GeneBank. The sequence collections are described in Table S2.

### Method to Detect Alternative Splicing

The process and software parameters used to assemble transcripts from Illumina short reads was described in full details (Mei et al. In Review). Briefly, three approaches were taken to assemble transcripts from Illumina short reads to maximize isoform detection. The assembly software used was annotation guided Cufflinks 2.2.1 (Trapnell et al. 2012), genome-guided Trinity 2.0.4 (Grabherr et al. 2011) and annotation guided StringTie 1.0.0 (Pertea et al. 2015). Isoforms built from cufflinks were required to have minimal expression FPKM value of 0.1 and required full reads support for the whole isoforms.

We masked vector and contaminant sequence in the Sanger ESTs using SeqClean (http://seqclean.sourceforge.net/). 454 transcripts were constructed using Newbler v2.8 (Roche 2010) with parameters “-cdna -urt”. Cufflinks, Trinity and StringTie assembled isoforms were merged with the Newbler 454 assemblies and the cleaned ESTs. Each species specific sequence collection was aligned back to the appropriate reference genome sequence followed by additional assembly and clustering to remove redundancy using the PASA 2.0 package (Haas et al. 2003). Isoform quality control steps described in Mei et al. (In Review) were used on the PASA outputted transcript set to remove isoforms that were poorly supported and/or potentially poor quality/poorly assembled. Briefly, each splice junction in a final retained isoform was required to have a minimum entropy score of 2 (Sturgill et al. 2013). Retained introns that define potential intron-retention isoforms were expected over a minimum of 90% of their length with a minimal median coverage depth of 10, and a minimum intron retention isoform ratio of 10%. The intron retention rate was calculated as described in Marquez et al. (2012), where the median coverage of the retained intron was divided by the number of reads supporting the splice junction. Lastly, we required each isoform to have a minimum FPKM of 1 and compose at least 5% of the total isoform abundance for that gene in at least one tissue examined.

We performed the above methods to identify and define AS events in rice, foxtail millet, *Brachypodium*, banana, African oil palm, and *Amborella*. The transcripts from Araport11 PreRelease 20151202 for *Arabidopsis* were realigned to the *Arabidopsis* reference genome using PASA to identify and characterize the set of *Arabidopsis* AS events. AS isoforms defined previously within maize and sorghum Mei et al. (In Review) were used to complete the AS isoforms sets analyzed in this study.

### Orthofinder Clustering Across Nine Species

Orthologous sequences were defined and grouped in Orthogroups across the nine species by OrthoFinder with default setting (Emms and Kelly 2015). OrthoFinder was designed for plant genome orthogroup identification and accounts for gene length bias.

### Identify Sets of Syntenic Genes Across Five Grass Species

Syntenic orthologs across five grass species (*Zea mays, Sorghum bicolor, Setaria italica, Oryza sativa, Brachypodium distachyon*) were defined using the methodology employed by (Schnable, Freeling, and Lyons 2012). Gene models were updated according to new annotations from Phytozome 10 (Goodstein et al. 2012). 11,995 syntenic gene lists across five species (sorghum, rice, foxtail millet, *Brachypodium*, either maize1/maize2 or both) were used for downstream analysis.

### Identification of Conserved Alternative Splicing

AS was independently identified in each species (see above) and all AS isoform sequences of the same event type (intron retention, exon skipping etc.) for a given species were pooled. The computational methodology to detect conserved AS relies on the creation of splice anchor sequence tags (SAST) for each AS event and then comparing the SASTs for a given event type with the SASTs defining the same event type between species. SASTs were generated for each AS event by extracting up to 300bp of sequence from both sides of the splice junction that defines the AS event types described in Figure 1. The tags from a particular AS event (i.e. exon skipping SASTs) from each species were compared to one another using tBLASTx from WU-BLAST (Gish, W. (1996-2003) http://blast.wustl.edu) with an E-value cutoff of 1e-5. If the length of two matching tags were both less than 30bp, this matching was not considered. In the cases tags being compared were less that 100 bp, the default tBLASTx E-value cutoff value was used. To classify conserved AS, SASTs flanking a AS event splice junction are required to be conserved between genes across species, and those genes must belong to the same orthogroup. Conserved AS events identified by this process were clustered and each cluster may contain conserved AS events identified between orthologous genes across species (1:1) as well as between paralogs. In an ideal setting, one conserved AS cluster represents one ancient conserved AS event across multiple species. However, if a gene within sorghum is present in two copies (homeologues) in maize, and all three genes harbor a conserved AS event, all three events will be assigned to the same event cluster. Therefore, with respect to the gene in sorghum and the two homeologous copies in maize, this cluster identifies 1 conserved event defined by 2 homeologous events in maize and 1 event in sorghum. Therefore, the total number of events making up a cluster may be larger than the number of species harboring the event. Because we identify these individual events within each species as instances of a conserved event we generalize the AS event cluster as a “conserved AS event”.

**Figure 1.**
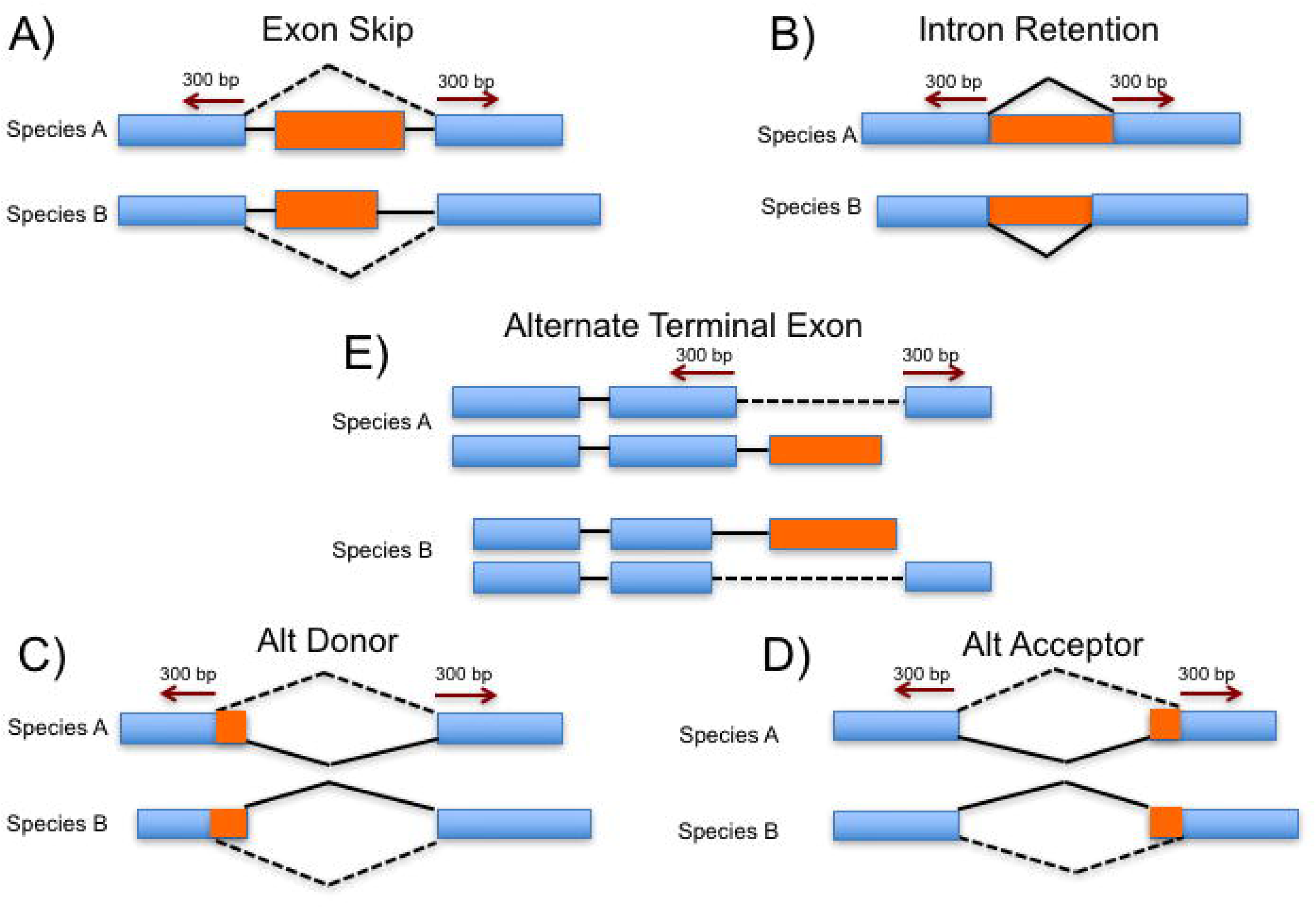
Pipeline to identify conserved alternative splicing events. Up to 300bp on either side of a splice junction were extracted. These extracted sequences were binned together with all other similarly extracted and binned sequences flanking the same type of event for each species (eg. one bin with sequences flanking junctions involved in intron retention, one bin with sequences flanking junctions involved in exon skipping, etc.). Sequences within each bin were compared with tBLASTx to bins of the same evet from other species to identify similar tags. Cases were the sequence tags were similar and the parent genes are from different species but represent orthologs as evidenced by being within the same orthogroup (see Methods) defined conserved alternative splicing events.

### Measure Selection Pressure on Alternative Splicing

We looked for evidence of selection pressure on AS events that were conserved by examining the difference between pairwise non-synonymous substitution rates (Ka) vs. synonymous substitution rates (Ks) for residues defined by exon sequences flanking the conserved AS event in grass species. These are the same regions that were used to construct SASTs (Figure 1). Both upstream and downstream alignments required minimal alignment lengths of 40 codons. Selection pressure on AS events at the amino acid levels was measured by determining the Ka/Ks ratios using KaKs_Calculator2.0 through model averaging (Wang et al. 2010). To minimize errors introduced by alignments, or due to paralogs that may result when comparing loci across large phylogenetic distances, the Ka/Ks evaluation of conserved AS events was limited to those events originating within orthologous grass genome loci that also exhibit conserved synteny (defined above).

### Testing the Functional Sharing Hypothesis: Whether Single Copy Genes Have More Conserved AS Events

The functional sharing hypothesis predicts that conserved AS events would have a higher incidence in single copy genes compared to non-single copy genes. We used a set composed of 2,717 multi-exon genes identified as “strictly” and “mostly” single copy identified across 20 flowering plant genomes by De Smet et al. (2013). We compared the number of single-copy genes exhibiting conserved AS events to the number of non-single copy genes with conserved AS events.

### Conserved AS in SR and hnRNP Proteins

Maize SR protein genes were retrieved from Rauch et al. (2014); Sorghum SR protein genes were retrieved from Richardson et al. (2011). Maize hnRNP protein encoding genes were retrieved from PlantGDB (Duvick et al. 2008). The primary protein sequences of each SR protein gene in maize and sorghum were used for phylogeny analysis. Protein sequences were aligned with MUSCLE (version 3.8.31) using default parameters (Edgar 2004). The alignments were used to construct a maximum likelihood phylogenetic tree with RAxML (version 8.1.12) software using a gamma distribution and LG4X model (Stamatakis 2014, Le, Dang, and Gascuel 2012). A manual search along the phylogenetic tree was performed to determine conserved AS events in SR families between maize and sorghum based on conserved isoform structure. The exon-intron gene structure was visualized with Fancygene version 1.4 (Rambaldi and Ciccarelli 2009) with AS events further labeled.

### Network Analysis in Stress Responsive Transcription Factors and Conserved Splicing Genes

3,150 *Arabidopsis* transcription factors (TFs) responsive to 14 diverse stresses were retrieved from STIFDB V2.0 (Naika et al. 2013) and we were focused on five major stresses (ABA, drought, cold, NaCl and light). Protein-protein interaction networks based on transcription factors (TFs) in *Arabidopsis* with conserved AS in response to five major stresses were built with using STRING v10 (Szklarczyk et al. 2014). In addition, a network based on *Arabidopsis* genes demonstrating conserved AS in *Amborella, Arabidopsis* and at least one monocot species was also constructed. The cluster of networks constructed was identified and analyzed in Cytoscape v3.3.0 (Shannon et al. 2003).

### Go Term Annotation

Genes harboring conserved AS events between *Amborella, Arabidopsis*, and at least one monocot species were further evaluated for functional annotation and GO term enrichment based on the *Arabidopsis* gene name using the agriGO analysis toolkit (Du et al. 2010). We use the Fisher’s exact test in singular enrichment analysis, followed by multiple test corrections with the Hochberg FDR test. Only associations with a minimum corrected p-value of 0.05 were considered.

### Data Availability

AS events table and GFF files for the AS isoforms in seven species (*Setaria italica, Oryza sativa L. ssp. Japonica, Brachypodium distachyon, Musa acuminate*, and *Elaeis guineensis*, the eudicot *Arabidopsis thaliana* and the basal angiosperm *Amborella*) are made available to the community via figshare (figureshare address will made immediately available once the manuscript has been accepted). In addition, a table detailing the AS events identified as conserved during this study will be made available. The *Zea mays* and *Sorghum bicolor* are already available community via figshare (http://10.6084/m9.figshare.4205079).

## Results

### Up to 54.6% of Expressed Multi-Exon Genes Exhibit Evidence of AS Among Nine Species

Five types of AS events were considered during this study: Intron retention (IntronR), Alternative acceptor (AltA), Alternative donor (AltD), Exon skipping (ExonS) and Alternate terminal exon (AltTE) (Figure 1). The percentage of expressed multi-exon genes that exhibit AS was determined for each species. The nine species evaluated exhibit AS across a range of 28.7% to 54.6% of multi-exon genes (Table S3). The largest proportion of expressed multi-exon genes with evidence of AS was in *Amborella* (54.6%). Over 60% of the intron retention events in *Amborella* were removed during the filtering process (described in Materials and Methods). Although the number of *Amborella* genes that exhibit AS is similar to that of other species examined during this analysis, *Amborella* has a lower proportion of multi-exon genes in the genome relative to other species. Therefore, despite the similar number of multi-exon genes that undergo AS, *Amborella* exhibits a high fraction of multi-exon genes that produce AS transcripts compared to the other species considered here. Intriguingly, *Amborella* genes that undergo AS produce the highest average number of isoforms/gene compared to other species (Table S3). Whether this feature reflects the evolutionary position of *Amborella* in angiosperm, or is the result of the absence of lineage specific whole genome duplication events – whereas species with additional lineage specific WGD events have lost or partitioned AS isoforms among paralogues through sub-functionalization (Jiang et al. 2013) – remains to be investigated. Maize has the largest collection of RNA-Seq data among the nine species examined and also has the largest number of AS events, but the percentage of multi-exon genes that undergo AS is similar to other species. While the proportions of genes that undergo AS and the total number of AS events is higher than previously reported for many plant species (Amborella Genome Project 2013, Campbell et al. 2006, Panahi et al. 2014, Abdel-Ghany et al. 2016, Walters et al. 2013), a substantially lower fraction of plant multi-exon genes undergo AS relative to human genes where a reported 95% of multi-exon genes undergo AS (Pan et al. 2008). Similar to previous genome-wide AS studies (Thatcher et al. 2014, Li, Xiao, and Zhu 2014, Mandadi and Scholthof 2015, Chamala et al. 2015), intron retention was the most commonly observed AS event type in monocots. Sorghum and *Amborella* were found to have the lowest percentage of intron retention events, 35.6% and 37.2%, respectively, while African oil palm and *Brachypodium* have the highest proportion of intron retention events, 61.4% and 57.6%, respectively (Table S3).

### AS Events Conserved Between Monocots, Arabidopsis and Amborella

We identified 9,804 conserved AS event clusters in nine species (Table 1). 59.0% of conserved AS events represent intron retention. Approximately 68.7%, 18.6%, 7.3% and 5.4% conserved AS events are conserved across two species, three species, four species and at least five species, respectively (Table 1). 1,816 conserved AS event clusters were found to include *Amborella*. Of these, 1,015 involved conserved intron retention events, 450 alternative acceptor events, 205 alternative donor events, 114 exon skipping events and 32 alternative terminal exon events (Table 2). In addition, we identified 80 conserved AS events conserved only between *Amborella* and *Arabidopsis* and absent from seven monocot species examined. These 80 AS events likely represent AS events lost after the divergence of eudicots and monocots including 34 intron retention, 31 alternative acceptor, 11 alternative donor, and 4 exon skipping events (Table 2). Analysis of nine species revealed 9,804 conserved AS event belong to 19,235 genes (Figure 2), with an apparent increased number of genes with conserved AS events within grass species (Figure 2); *Arabidopsis* had the least (944) number of genes harboring conserved AS events while maize had the most (3,687).

**Figure 2.**
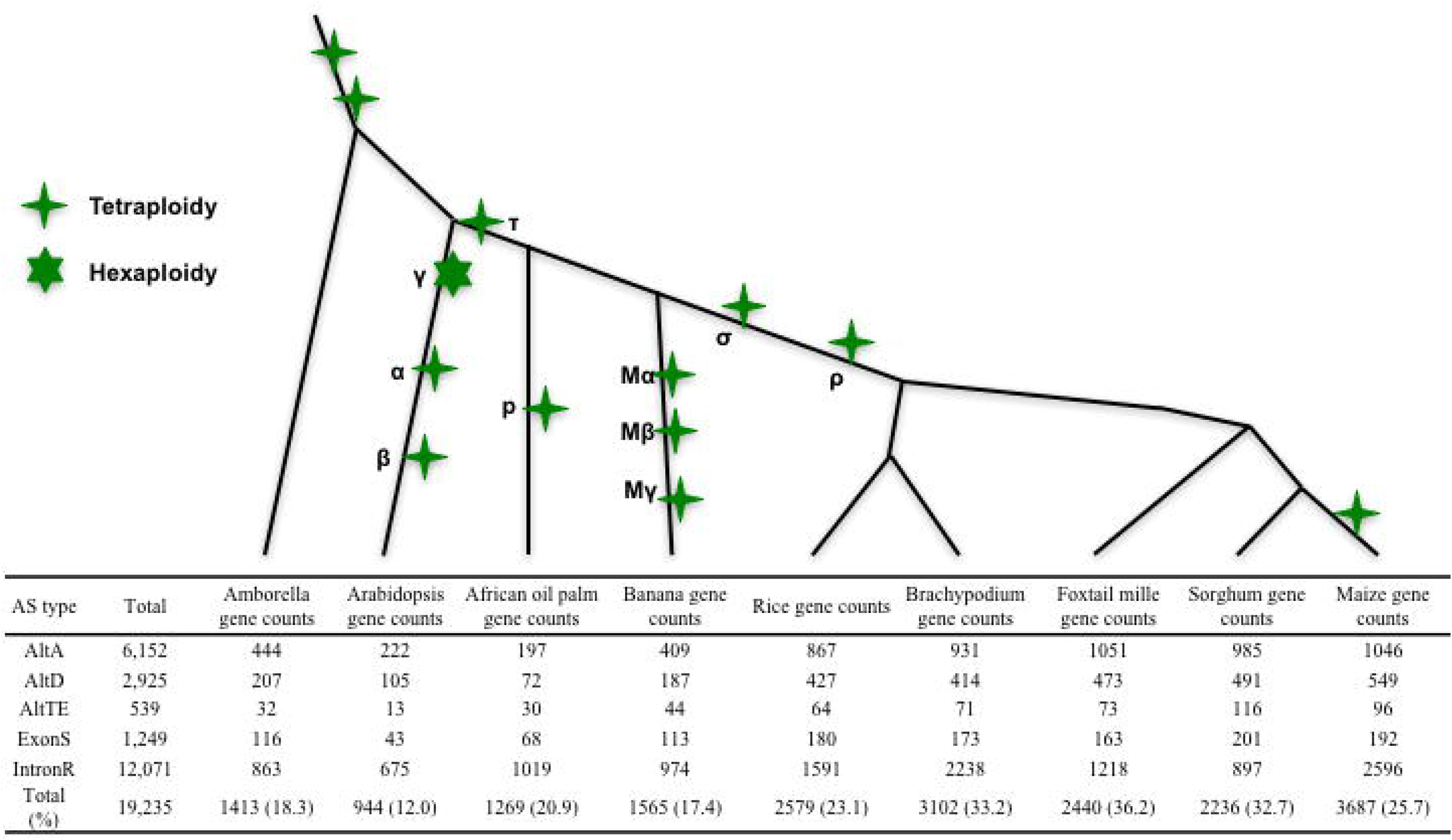
Genes with conserved alternative splicing events across nine species. We calculate the number of genes with conserved alternative splicing events in each species. Tetraploidy and Hexaploidy are labeled on the phylogenetic trees. The length of the phylogenetic tree is not proportional to the phylogenetic distance.

**Table 1.** Conserved alternative splicing across 9 species at the gene family level. Percentage is based on total number of conserved alternative splicing clusters.

| Number of species | AltA | AltD | ATE | ExonS | IntronR | Total/% |
| --- | --- | --- | --- | --- | --- | --- |
| 2 | 1,384 | 684 | 201 | 335 | 4,128 | 6,732/68.7 |
| 3 | 447 | 181 | 32 | 69 | 1,099 | 1828/18.6 |
| 4 | 218 | 112 | 6 | 26 | 351 | 713/7.3 |
| 5 | 117 | 36 | 2 | 21 | 109 | 285/2.9 |
| 6 | 58 | 29 | 0 | 10 | 52 | 149/1.5 |
| 7 | 22 | 12 | 0 | 3 | 32 | 69/0.7 |
| 8 | 7 | 3 | 0 | 4 | 7 | 21/0.2 |
| 9 | 2 | 2 | 0 | 1 | 2 | 7/0.1 |
| Total/% | 2,255/23.0 | 1,059/10.8 | 241/2.5 | 469/4.8 | 5,780/59.0 | 9,804 |

**Table 2.** Conserved alternative splicing clusters among different species and clades.

\* Conserved at least one species in PCAMAD clade and one species in BEP clade.
|  | Total | IntronR | AltA | AltD | ExonS | ATE |
| --- | --- | --- | --- | --- | --- | --- |
| Conserved between *Amborella* and other species | 1,816 | 1,015 | 450 | 205 | 114 | 32 |
| Conserved in *Amborella*, *Arabidopsis*, and at least one species of Monocots | 129 | 64 | 43 | 11 | 10 | 1 |
| Conserved between *Amborella* and *Arabidopsis* not in Monocots | 80 | 34 | 31 | 11 | 4 | 0 |
| Conserved across *Amborella* and seven Monocot not in *Arabidopsis* | 9 | 2 | 3 | 2 | 2 | 0 |
| Conserved in banana, African oil plum and at least one grass species | 239 | 162 | 44 | 20 | 11 | 2 |
| Conserved across five species in grass | 224 | 60 | 106 | 37 | 21 | 0 |
| Conserved across seven species in monocots | 34 | 13 | 12 | 5 | 4 | 0 |
| Conserved between PACMAD and BEP clades * | 4333 | 2548 | 1052 | 480 | 177 | 76 |

GO term enrichment analysis was performed on *Arabidopsis* genes with AS events that are conserved within *Amborella, Arabidopsis* and monocots to identify whether genes with conserved AS events participated in particular biological processes. 129 AS events meet this criteria. These 129 conserved AS events represent 64 intron retention events, 43 alternative acceptor events, 11 alternative donor events, 10 exon skipping events and 1 alternative terminal exon events (Table 2). GO term enrichment based on the *Arabidopsis* genes producing these 129 conserved AS events suggest an over-representation of kinase and phosphorylation activity (Figure S1). This result is consistent with the proposal that AS impacts protein kinase mediated signaling pathways (Reddy et al. 2013), which may modulate the transfer of developmental or environmental cues.

TFIIIA possesses an exon skipping event conserved across land plants (Fu et al. 2009, Barbazuk 2010). This event was also identified in all species examined in our study, although our stringent depth requirements initially filtered it out of the banana analysis. In addition, a widely conserved AS event was discovered within SPA3, a member of the SPA protein family, that plays a pivotal role in light signaling as a suppressor of photomorphogenesis (Laubinger and Hoecker 2003). Initial investigation by Shikata et al. (2014) identified an intron retention event and an alternative donor event within *Arabidopsis* SPA3; both splice isoforms harbor premature terminal codons that result in truncated protein products. These truncated proteins retain interaction with CONSTITUTIVE PHOTOMORPHOGENIC 1 (COPI) but lose the ability to bind to DAMAGED DNA-BINDING PROTEIN 1 (DDB1). Evidence suggests the phytochrome could mediate the production of AS transcripts of SPA3 in response to some light conditions, thus promoting plant photomorphogenesis (Shikata et al. 2014). Both the intron retention and alternative donor AS events were identified in our analysis: the alternative donor event is conserved across all nine species examined and the intron retention event is conserved in four species: maize, foxtail millet, African oil palm and *Arabidopsis*. The absence of the intron retention event in the other species examined could be due to limited data depth or tissue sampling. Further investigation is required to determine whether or not this intron retention is conserved broadly across the angiosperms.

Alternative donor and acceptor events were reported to be significantly enriched among events conserved between *Brassica* and *Arabidopsis* (Darracq and Adams 2013). We found significant enrichment of alternative donor and alternative acceptor events conserved across at least three species relative to events conserved in only two species (P<0.0001, Fisher’s exact test, Table 1), similar enrichment patterns were observed for exon skipping events conserved across at least four species relative to clusters conserved in only two species (P<0.05, Fisher’s exact test, Table 1). Expectedly, the overall numbers of AS events that are conserved decreases as the number of species they are conserved within increases. In addition, when considering AS conserved across multiple species, as the number of species that harbor the conserved AS increase, the proportion of conserved AS that are intron retention decrease relative to the other event types. This suggests that requiring an AS event to be conserved between species selects for biologically relevant and important events, and that the trend we observe with intron retention might imply that intron retention suffers a higher proportion of ‘noisy’ or non-relevant splicing relative to other AS types.

Examining the lengths of conserved retained introns relative to the non-conserved retained introns across 9 species does not reveal a universal trend. In *Arabidopsis, Brachypodium*, and foxtail millet, the length of conserved retained introns is significantly shorter than that of non-conserved (i.e. unique to only one species) retained intron (Table 3). However, in *Amborella* and African oil palm, the length of conserved retained introns is significantly longer than that of non-conserved retained introns. In banana, maize, sorghum and rice, there is no significant difference in retained intron length between conserved and non-conserved retained introns. However in the related grass species maize, rice and sorghum that exhibit large segments of genome co-linearity (Gale and Devos 1998) the mean length of conserved retained intron is shorter than non-conserved retained introns, while the median lengths of conserved retained introns are similar or larger than those of non-conserved retained introns (Table 3). A previous report suggested that the lengths of retained introns within *Arabidopsis* that are conserved in *Brassica* are shorter than non-conserved retained introns (Darracq and Adams 2013). We observed the same length trend in *Arabidopsis*, but this is not consistent in other species.

**Table 3.** A comparison of intron length in the conserved intron retention events vs. non-conserved intron retention events across the species examined. (NS, Not significant; *, P < 0.05; ***, P < 0.001)

|  | *Amborella* | *Arabidopsis* | Banana | *Brachypodium* | Foxtail millet | Maize | Oil Palm | Rice | Sorghum |
| --- | --- | --- | --- | --- | --- | --- | --- | --- | --- |
| Conserved AS |  |  |  |  |  |  |  |  |  |
| Mean | 670 | 165 | 329 | 411 | 402 | 440 | 315 | 417 | 409 |
| Median | 390 | 96 | 143 | 233 | 237 | 207 | 131 | 269 | 256 |
| No evidence for conserved AS |  |  |  |  |  |  |  |  |  |
| Mean | 651 | 186 | 327 | 474 | 410 | 471 | 287 | 486 | 437 |
| Median | 316 | 113 | 131 | 245 | 201 | 205 | 121 | 247 | 242 |
| Wilcoxon test |  |  |  |  |  |  |  |  |  |
| P value | *** | *** | NS | *** | * | NS | * | NS | NS |

*Arabidopsis* TFs responsive to 14 diverse stress signals were retrieved from STIFDB V2.0 (Naika et al. 2013). TFs responsive to oxidative stress, dehydration, ABA, NaCl, drought, light, and cold have more AS relative to TFs responsive to the other 7 stresses (Table S4). Since the oxidative stress and dehydration responsive TFs only have 1 and 8 TFs with conserved AS, respectively, we focus on the five major stresses where AS appears to play a significant role: ABA, NaCl, drought, light, and cold. The absolute number and percentage of AS in these TFs responsive to five major stresses are described in Table S4 together with the number and the percentage of conserved AS in these TFs. Two genes are particularly interesting, AT4G27410 (NAC transcription factor RD26) and AT1G78070 (Transducin/WD40 repeat-like superfamily protein). These two genes have conserved AS events and are responsive to all five stresses based on the data (Figure S2). RD26 has a conserved intron retention event across 7 of the 9 species we examined (absent in sorghum and *Brachypodium*). AT1G78070 has a conserved alternative terminal exon between sorghum, banana and *Arabidopsis*. These results indicate that there are AS events in stress-response TFs conserved among species across large phylogenetic distances, which suggests that AS may play an important role in the activity of some TFs during stress response.

### AS Conserved Between Monocot Grass and Non-Grass Species

There are 239 AS events conserved between banana, African oil palm, and at least one of the five grass species examined (Table 2). 34 out of 239 conserved AS events are conserved across banana, African oil palm and five grass species (Table 2). 204 conserved AS events exist within the banana and African oil palm lineages but are absent from the grass species studied. 224 conserved AS events are conserved across the five grass species examined, and these are comprised of 60 intron retention events, 106 alternative acceptor events, 37 alternative donor events and 21 exon skipping events (Table 2). Within the PACMAD clade, 1,544, 1,488 and 1,323 AS events are conserved between maize vs. sorghum, maize vs. foxtail millet, and sorghum vs. foxtail millet, respectively. In the BEP clade, 1,658 conserved AS events are conserved between rice and *Brachypodium*. 4,333 conserved AS events are conserved in at least one species in the BEP clade and one species in the PACMAD clade (Table 2). Maize clock genes (i.e circadian rhythm) GRMZM2G033962 (pseudoresponse regulator protein 37, *PRR37*) and GRMZM2G095727 (pseudoresponse regulator protein 73, *PRR73*) share a conserved intron retention that is also conserved across rice, *Brachypodium*, and foxtail millet. Our data suggests that this intron retention event is likely specific to members of the grass family; the role of intron retention in *PRR37* and *PRR73* controlling photoperiodic flowering needs further experimental investigation.

Since syntenic genes were more likely to transcribe transcripts and translate proteins than non-syntenic regions (Walley et al. 2016), we tested the hypothesis that AS and conserved AS is enriched in syntenic regions across grass species compared to non-syntenic regions. There are 11,996 syntenic gene clusters across five grass species including maize, sorghum, foxtail millet, rice, and *Brachypodium*. Since maize has undergone an additional whole genome duplication relative to the others, syntenic genes could be maize1 gene, maize2 gene or both. In each of five grass species, we compared the proportion of genes within syntenic relationships that undergo AS vs. non-syntenic genes, and the proportion genes within syntenic relationships that have conserved AS events vs. the proportion of non-syntenic genes that have conserved AS events. The proportion of genes with syntenic relationships that exhibit AS is approximately twice that of genes in non-syntenic relationships (Figure 3). This trend is consistent across five grass species. Similarly, the proportion of genes that have conserved AS events that reside in syntenic regions is approximately twice that of genes with conserved AS events that do not reside within syntenic regions (Figure 3). This suggests that genes within conserved syntenic blocks between present within the five grass species studied are enriched in both AS, and conserved AS events.

**Figure 3.**
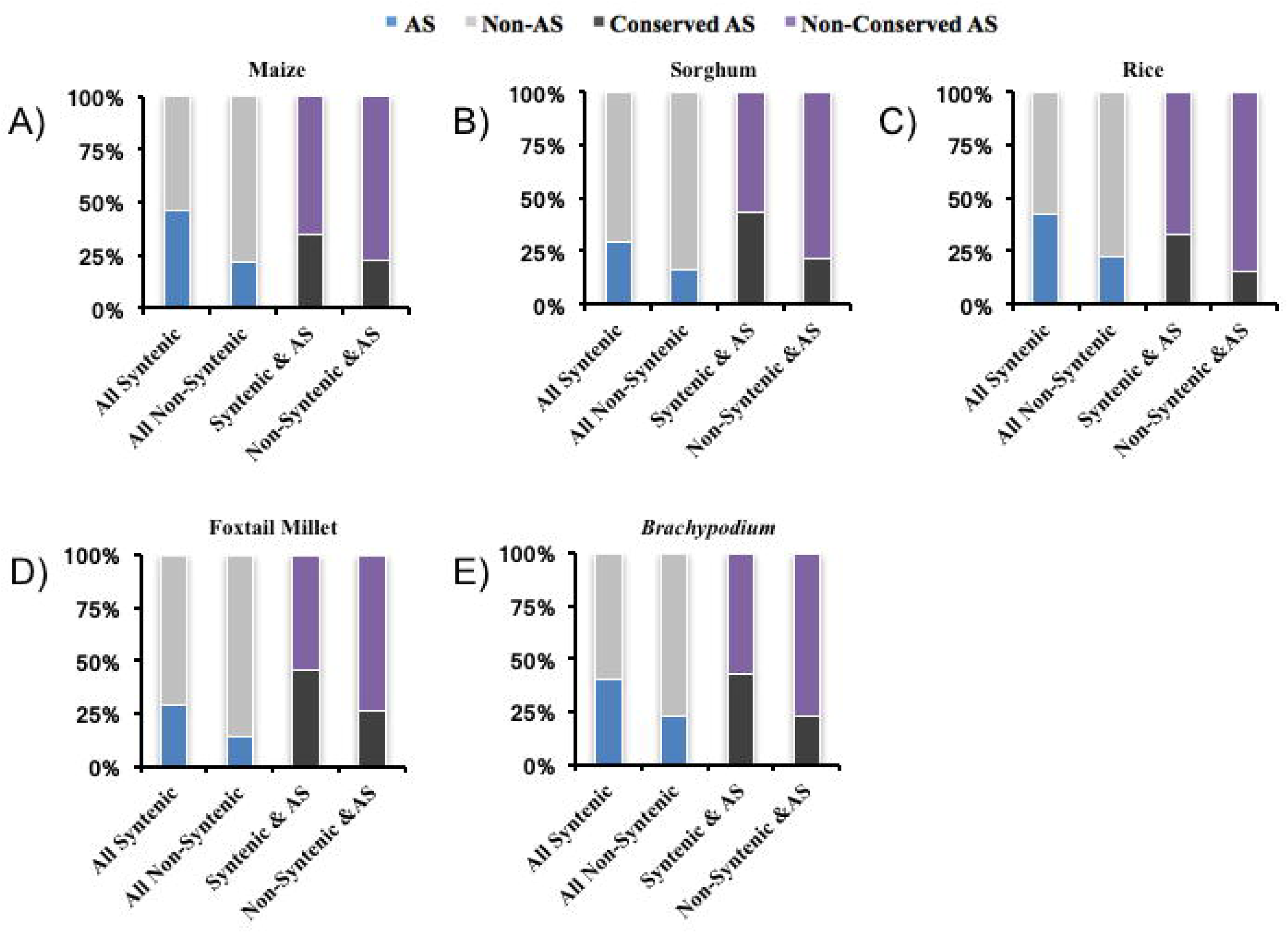
Alternative splicing and conserved alternative splicing enriched across grass syntenic genes. We calculated the percentage of AS in syntenic and non-syntenic genes (the percentage of AS genes are labeled in blue). In addition, we calculated the percentage of conserved AS in syntenic AS genes and non-syntenic AS genes (the percentage of conserved AS genes are labeled in black).

We examined selection pressure at the amino acid level on AS events conserved within grasses by performing pairwise comparisons of Ka/Ks ratios across the exonic regions flanking each conserved AS event (Figure 4). In total 4,083 flanking exonic regions were examined, 1,999 from upstream exons and 2,084 from downstream exons. We separated the Ka/Ks values for the flanking upstream and downstream portions of each alternative splicing type and plotted these values (Figure 4). In general, Ka/Ks values of exonic regions flanking conserved AS events are smaller than 1 (Figure 4), which suggest these regions are under purifying selection. There are 161 pairwise flanking region pairs with Ka/Ks value above 1, indicative potential candidate for positive selection. These include 67 from intron retention, 41from alternative acceptor, 40 from alternative donor, 8 from exon skipping and 5 from terminal alternate exon. We did not detect a statistically significant difference in Ka/Ks between exons upstream and downstream of conserved intron retention, alternative acceptor, and exon skipping events. However, a significant difference in Ka/Ks was observed between upstream and downstream exons flanking conserved alternative donor events (P < 0.001, Wilcoxon Rank-Sum Test) and alternate terminal exon (P < 0.001, Wilcoxon Rank-Sum Test). In general, these results suggest that the regions flanking conserved AS events in grasses are undergoing purifying selection, which may be maintaining *cis*-signals required for AS and somewhat responsible for maintaining sequence conservation across species.

**Figure 4.**
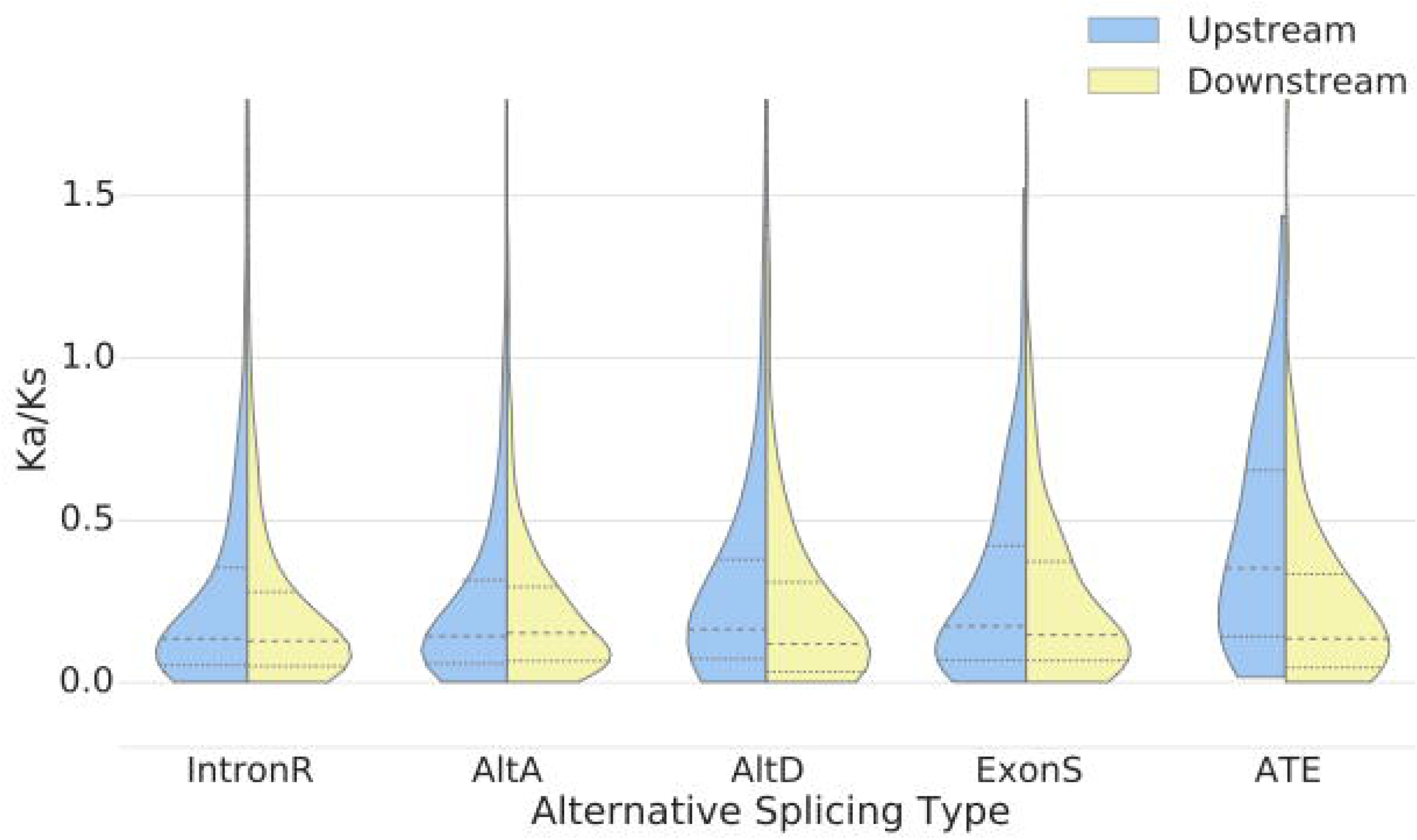
Violin plot of Ka/Ks value from flanking upstream and downstream in different alternative splicing events across syntenic genes within grass species. We calculated the pairwise Ka/Ks ratio within exonic regions upstream and downstream of conserved alternative splicing events in five grass species. Only those exonic regions aligned at a>= 120bp were considered in Ka/Ks calculation. The dotted lines within the violin plot represented the first, second and third quartile.

### Conserved AS is More Common Among Single Copy Genes

We tested the functional sharing hypothesis between single copy genes vs. non-single copy genes by measuring the number of genes within each category that exhibit conserved AS events. 2,985 single copy genes including strict and mostly single copy were described in *Arabidopsis* (De Smet et al. 2013). Han et al. (2014) suggested single copy genes had increased levels of AS relative to genes from large gene families and a decreased proportion of AS compared to genes from small gene families. Similarly, Su and Gu (2012) suggested that single copy genes had more isoforms per gene compared to the genes from large gene families but similar average isoform counts per gene compared to the genes from small gene families.

We compared the number of conserved AS genes between single copy genes vs. non-single copy genes irrespective of gene family size. 2,717 out of 2,985 single copy genes are multi-exon genes. 1,113 out of the 2,717 multi-exon genes have evidence of AS in *Arabidopsis* based on the Araport11 transcripts. In the 19,700 non-single copy multi-exon genes, there are 6,751 that exhibit AS. Among multi-exon genes, the single copy genes have significantly more AS compared to the rest of multi-exon genes (P < 0.0001, Chi-square with Yates’ correction). Among the 944 *Arabidopsis* genes with conserved AS events in our analysis, 173 were from 1,113 single copy AS genes, while the remaining 771 were from the 6,751 non-single copy AS genes. Single copy genes in *Arabidopsis* showed significantly enriched in conserved AS events (P < 0.0001, Chi-square with Yates’ correction). These results suggest that among multi-exon genes in *Arabidopsis*, a greater proportion of single copy genes both undergo AS and harbor conserved AS events than do non-single copy genes.

### Conserved AS is More Common Among SR Protein Genes Relative to hnRNP Proteins

Similar to previous observations (Kalyna et al. 2006, Rauch et al. 2014, Chamala et al. 2015), AS events were detected in SR proteins. 19 out of 21 SR proteins in maize demonstrate AS, and 13 out of 19 alternatively spliced SR protein genes in maize showed 24 conserved AS events with other species including 13 intron retention events, 4 alternative acceptor events, 4 alternative donor events and 3 exon skipping events (Table S5). 10 out of 24 conserved AS events are present in at least five species. Richardson et al. (2011) identified two SR proteins (GRMZM2G110143 and GRMZM2G170365) in maize with evidence of positive selection (Supplemental Table5). These two genes also have conserved AS events across 2 and 4 species, respectively. Zm-SCL30 (GRMZM2G065066), previously identified in maize, responds to both cold and heat response (Mei et al. In Review). SCL30 has alternative acceptor, alternative donor, and exon skipping events conserved across grass species and *Amborella*, as well as two intron retention events only conserved within the grass lineage. Altogether, there are 4 maize SR proteins (2 in the RS subfamily, 1 in the SC subfamily, and 1 in the SCL subfamily) that demonstrate 6 conserved AS events with *Amborella* (Table S5), which supports the existence of a deeply conserved splicing mechanism in SR proteins. 10 out of 40 hnRNP proteins in maize exhibit evidence for 17 conserved AS events, 10 of which are intron retention events, 4 alternative acceptor events, 2 alternative donor events, and 1 exon skipping events (Supplemental Table5). None of these 17 conserved AS events are conserved across more than four species. 3 of the 17 conserved AS events from 3 hnRNP genes in maize are conserved with *Amborella* (Table S5), while 6 out of 24 AS events from 4 SR proteins in maize are conserved with *Amborella*. These results suggest that the signals for AS within SR protein genes across species are better conserved than those within the hnRNP protein encoding genes.

### Conservation and Subfunctionalization of SR Proteins After Whole Genome Duplication

We used a phylogenetic approach to identify conserved AS across the SR protein subfamilies in maize and sorghum. An ancestral exon skipping event is conserved across all members of this subfamily in maize and sorghum (Figure 5A). Intriguingly, all three maize SR subfamily genes were reduced to one copy after whole genome duplication; one gene lost a homeologous copy from maize subgenome1, and two genes lost their homeologous copies from subgenome2. The plant-specific RS subfamily also has a shared exon skipping in the second long intron region coding RRM domain both in maize and sorghum (Figure 5A). The exon skipping isoform generates a complete RRM domain. Kalyna et al. (2006) suggested this long intron is conserved from green algae to angiosperms, and the splice site is retained between monocot and eudicot lineages. Our analysis suggests that this exon skipping event is preserved in maize, sorghum, rice, *Brachypodium*, foxtail millet and *Amborella*. In the plant-specific RS2Z subfamily, maize maintained both copies of the genes after whole genome duplication. Remarkably, two sorghum genes Sobic.009G022100 and Sobic.009G022200 are next to each other but in opposite directions on the chromosome, which suggests one copy may be the result of a tandem duplication. Based on phylogenetic evidence Sobic.009G022200 might be the older copy (Richardson et al. 2011). Not only did Sobic.009G022200 and two duplicate genes in maize retain an exon skipping event, but they also contain two intron retention events surrounding the skipped exon (Figure 5B). Sobic.009G022100 and the two duplicate genes in maize (GRMZM2G099317 and GRMZM2G474658) apparently diverged their AS isoforms (Figure 5B). Sobic.009G022100 only preserved the exon skipping isoform, which would generate a complete RRM domain. GRMZM2G099317 retained the intron retention event on the right side, and the exon skipping event. For the second copy, GRMZM2G474658, the intron retention event on the left side is retained as well as an alternative acceptor event. There is one additional sorghum gene Sobic.003G064400 without conserved synteny in maize, which suggests maize might have lost both syntenic copies that had represented orthologues of this sorghum gene. The AS pattern of this gene is similar to AS pattern of GRMZM2G474658 (Figure 5B). The alternative long intron in the RS2Z subfamily is also conserved from mosses to angiosperms (Kalyna et al. 2006).

**Figure 5.**
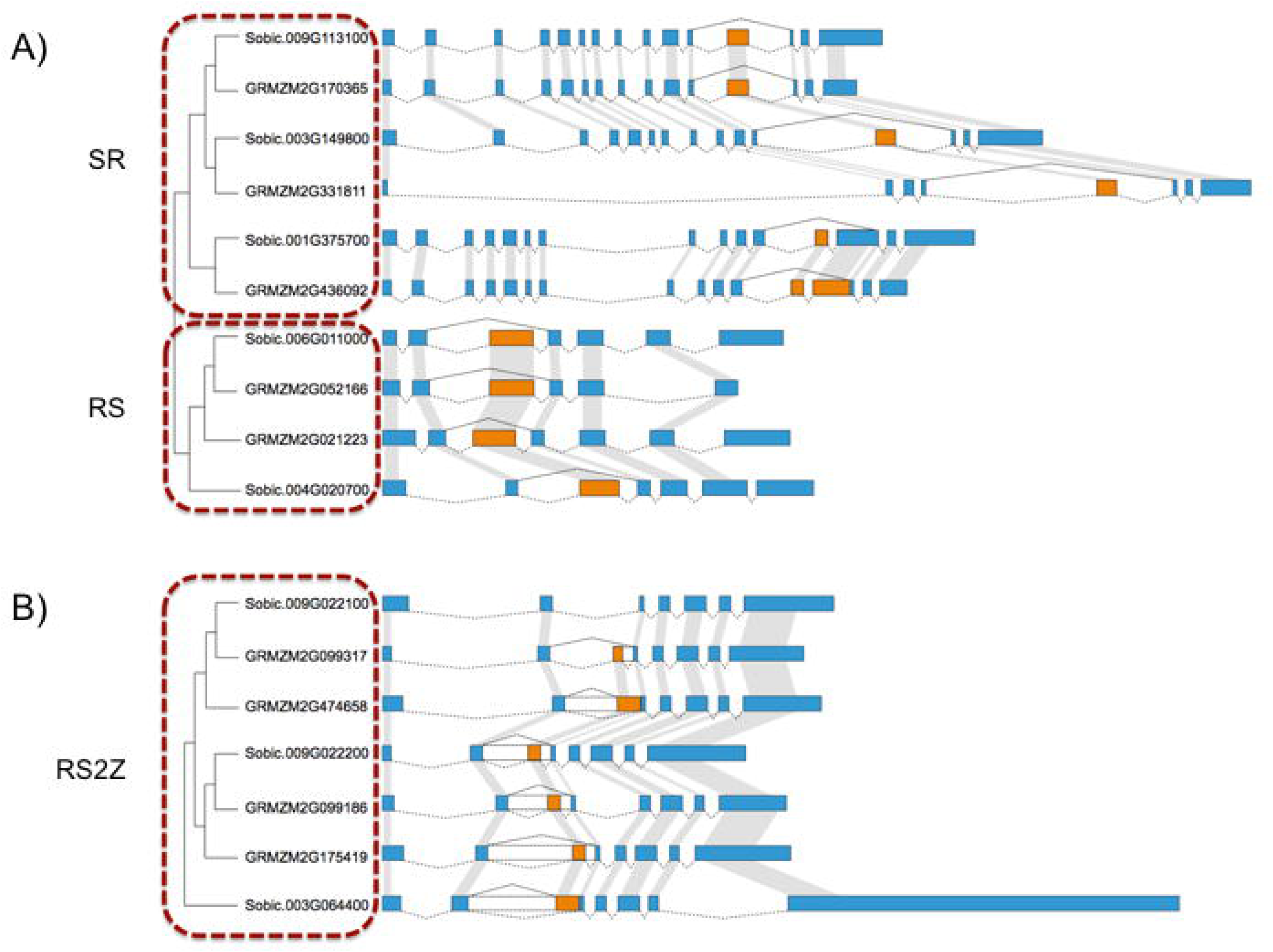
Phylogenetic tree and gene structure of SR, RS and RS2Z subfamilies in maize and sorghum. A) Phylogenetic tree representing the relationship of genes from SR and RS subfamilies in maize and sorghum; associated gene structures are illustrated by Fancygene to the right. Constitutive exons are shaded blue and alternative region are indicated with orange. Transcript structure is indicated by exons connected via dashed lines; orthologous exons are indicated with grey lines. B) Phylogenetic tree and gene structure of RS2Z subfamily in maize and sorghum are illustrated by Fancygene. Annotation and shading is described above; intron retention regions are indicated in clear boxes.

### R2R3-MYB is Tightly Connected with a Plant-Specific SR Protein Subfamily

We used a network building approach to identify the degree of interactivity of TFs with conserved AS in response to five stresses (ABA, drought, cold, NaCl and light) based on protein-protein interactions in STRING v10 (Szklarczyk et al. 2014). Several interesting relationships showed up in the network (Figure S3). Ubiquitin genes such as UBQ10, UBQ11, and UBQ14 are centralized in the network, which suggests AS might play a role in the susceptibility of translated isoforms to ubiquitin mediated degradation. Ubiquitin genes are closely associated with splicing related genes (such as RS2Z33, RS41, RS2Z32, RS40, U2AF65A, SR45, SCL33) via *Glycine Rich Protein 7* (GRP7). GRP7 is part of the circadian clock and negatively auto-regulates its own protein abundance by producing a non-productive AS isoform that is subject to the nonsense-mediated decay (NMD) pathway (Schöning et al. 2007, Schöning et al. 2008). In addition, we identified the MYB gene network is associated with splicing network genes through the proline-rich spliceosome-associated protein (AT4G21660) (Figure S3). The MYB genes identified in this network belong to the R2R3-MYB family. The R2R3-MYB family has undergone expansion in plants (Du et al. 2014) and plays many important plant-specific roles. Furthermore, there are many kinase and phosphatase related genes in close association with genes responsible for phosphorylation and dephosphorylation of splicing regulators, such as serine-threonine protein kinase (CIPK3), calcium-dependent protein kinase 29 (CPK29), CBL-interacting protein kinase 9 (CIPK9), and calcineurin b-like protein (CBL1). CBL1 has been identified as a salt-tolerance gene and connects with components of the spliceosome (Feng et al. 2015). CBL2 and CBL3 together with CIPK3 and CIPK9 can form a calcium network to regulate magnesium levels, while CBL1 is a calcium sensor (Tang et al. 2015). Here, AS of CBL1 and CIPK3, CIPK9 could potentially participate in regulating magnesium levels through a feedback loop sensitive to calcium concentration.

We also examined the networks containing genes in *Arabidopsis* with conserved AS events with *Amborella* and at least one species of monocot. Most of the genes present in the network were from the plant-specific SR protein subfamily, such as RS2Z, RS, and SR45a. In addition, TFIIIA is connected to the splicing protein network via RNA-binding protein (AT4G35785). TFIIIA has an exon skipping event that is conserved across the land plants (Barbazuk 2010, Fu et al. 2009). Similar to the network based on TFs in response to stress, we identified a close connection between the R2R3-MYB class with a splicing protein network via TFIID-1 (AT3G13445) and proline-rich spliceosome-associated family protein (AT4G21660) (Figure 6). These R2R3-MYBs are involved in many important functions, such as glucosinolate biosynthesis, phenylpropanoid pathway, conical epidermal cell outgrowth, drought and pathogens ABA- and JA-mediated, and cuticular wax biosynthesis (Table S6).

**Figure 6.**
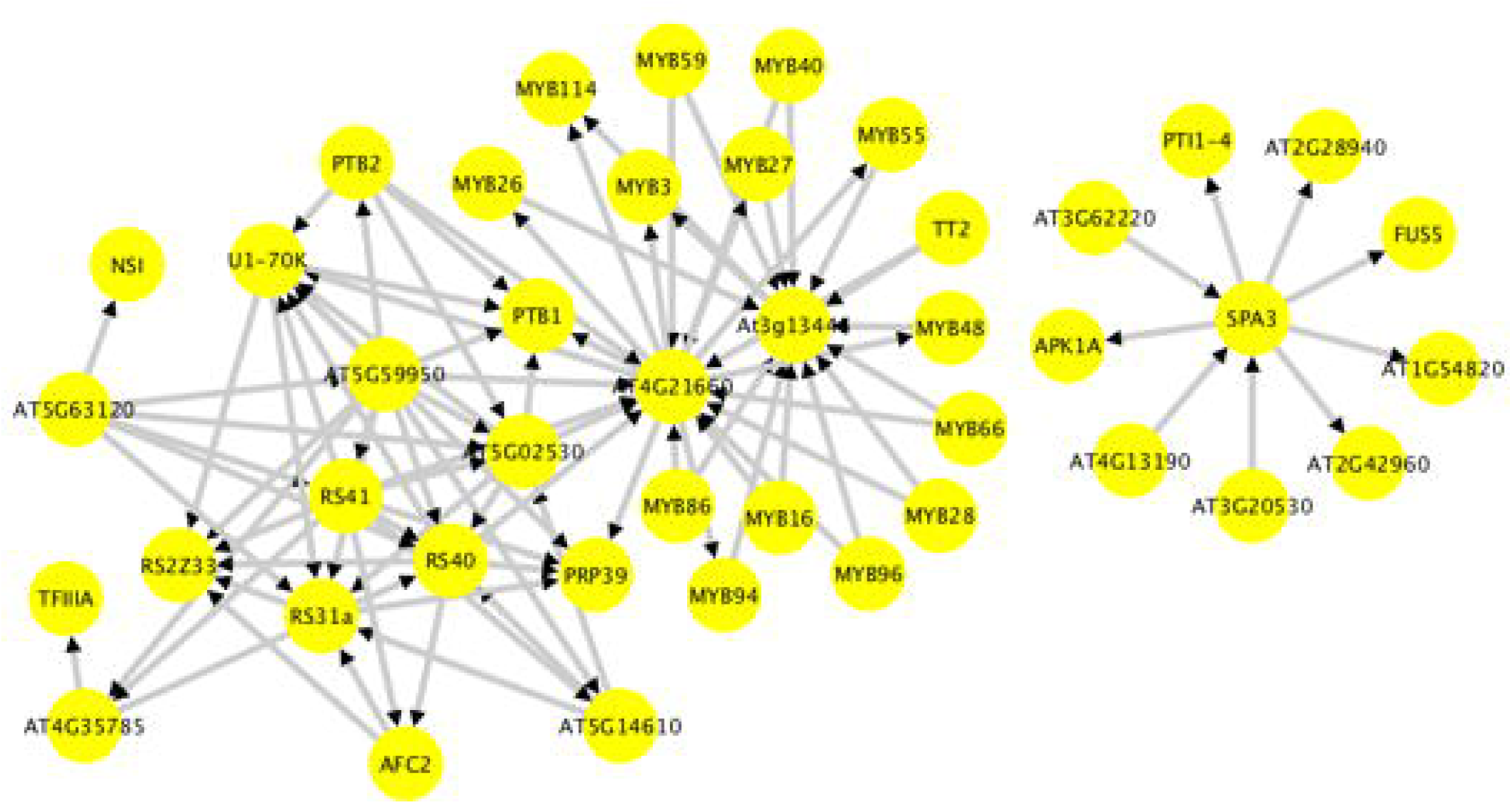
Protein-protein network of *Arabidopsis* genes with evidence of conserved alternative splicing with *Amborella* and at least one species of monocot. The arrow in the network indicates the direction of the protein-protein network.

## Discussion

In this study, we utilized a bioinformatics scheme to detect AS events and identify those which are conserved across 9 plant species by mining publicly available RNA-Seq data. In total, across all 9 species, 9,804 AS events conserved between 2 or more species are defined within 19,235 genes. New sequenced plant genomes and vast amounts of RNA-Seq are increasingly deposited into NCBI’s Sequence Read Archive (SRA). These are treasure troves of data waiting to be mined for new discoveries. Many are already tapping into this wealth, as exemplified by (Nellore et al. 2016) who examined splice junction variants in the human genome, and Chamala et al. (2015) who examined conserved AS events across several species of eudicots. The method we describe here can be used to mine to identify conserved AS events within any species with an available genome sequence and deep RNA-Seq data.

### AS Shows Widely Conserved Across Species but Likely Undergone Complicated Patterns of Gain and Loss

Each species examined has one to three thousand genes with evidence of conserved AS. *Arabidopsis* has least number of genes with conserved AS (944), while maize has the greatest number of genes (3,687) (Figure 2). The smallest percentage of conserved AS genes among genes that undergo AS is *Arabidopsis* (12.0%), while foxtail millet has the greatest percentage of conserved AS genes among genes with AS (36.2%). We identified 1,413 genes with conserved AS events in *Amborella*, which account for 18.3% of the genes in *Amborella* with evidence of AS. One conserved exon skipping event occurs within the gene encoding an RNA-binding KH domain-containing protein. This conserved AS event was detected in 8 of the 9 species RNA-Seq data examined, but was apparently absent in *Arabidopsis*. However, this isoform is reported within the AtRTD *Arabdiopsis* transcriptome data (Zhang et al. 2015) indicating that it is broadly conserved across all species investigated here. RNA-binding KH domain proteins are vital for heat stress-responsive gene regulation (Guan et al. 2013). The alternatively spliced exon within this RNA-binding KH domain-containing protein has high sequence similarity across nine species studied (Figure 7) and this AS event is likely conserved broadly across angiosperms. Similar to a previous study that identified that alternative donor and alternative acceptor events were found to be significantly enriched between *Arabidopsis* and *Brassica* (Darracq and Adams 2013), we identified a significant enrichment of alternative donor and alternative acceptor events that were conserved in at least three species relative to those conserved in only two species, and a significant enrichment of exon skipping events conserved in at least four species relative to those conserved in only two species. We observed that *Arabidopsis, Brachypodium* and foxtail millet retained introns that are conserved have significantly shorter lengths compared to non-conserved retained introns, which is consistent with a previous report in *Arabidopsis* (Table 3) (Darracq and Adams 2013). However, this trend is not observed across all the species examined. Conserved retained introns in *Amborella* and African oil palm are longer than those retained introns that were not conserved, and there are no significant differences in the lengths of conserved retained introns vs. non-conserved introns in banana, maize, rice, and sorghum.

**Figure 7.**
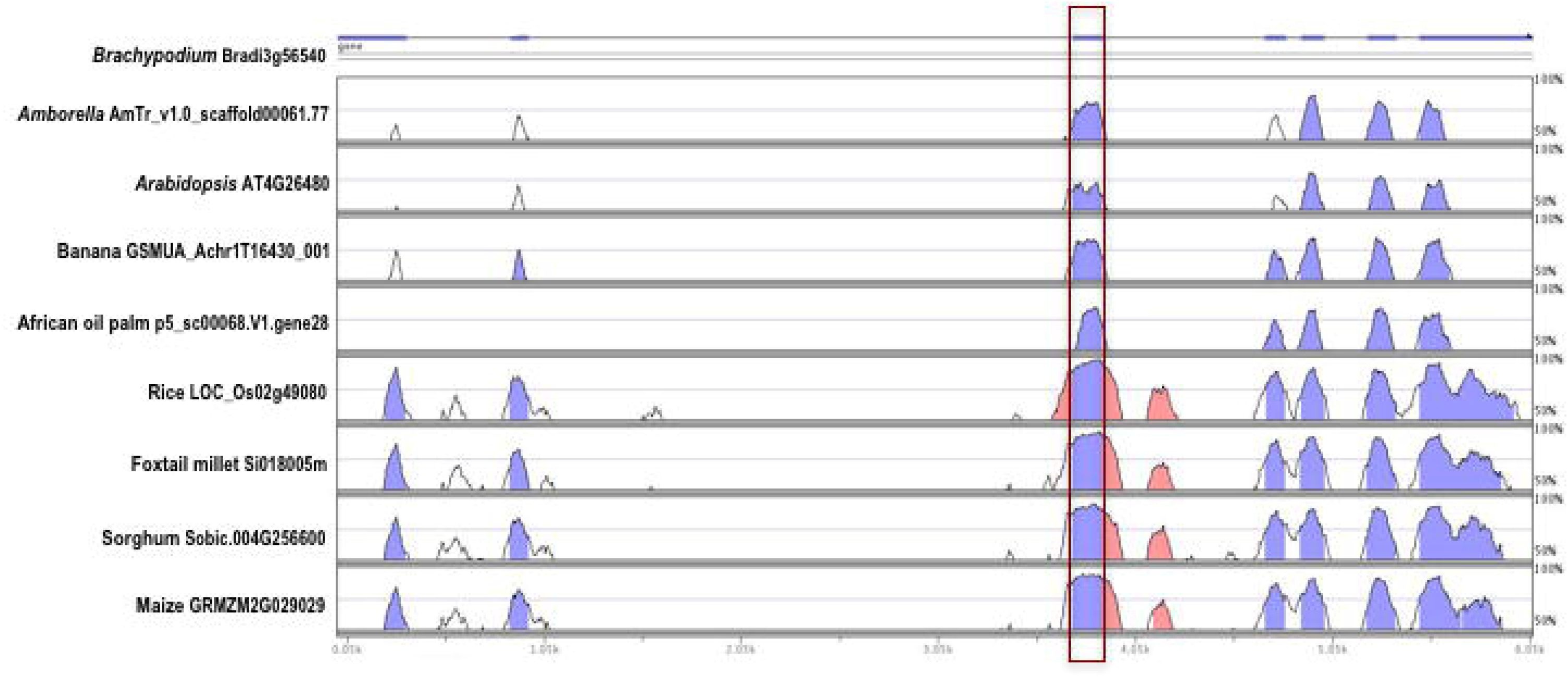
Sequence conservation within a gene encoding an RNA-binding KH domain-containing protein that produces an exon skipping even conserved across species. An example of sequence conservation of the alternative exons in a RNA-binding KH domain-containing protein. The exon skipping in *Arabidopsis* is not included in the Araport11, but detected in AtRTD. Vista plots of pair-wise alignments between the RNA-binding KH domain protein encoding gene from *Brachypodium* and its orthologue in *Amborella, Arabidopsis*, banana, African oil palm, rice, foxtail millet, sorghum and maize. The alternative exon (boxed in red) is highly conserved between *Brachypodium* and its orthologues in eight other plant species. Levels of sequence identity between *Brachypodium* and its orthologues in eight other plants species displayed are depicted as blue peaks. Pink bars signify segments that pass the alignment criteria of 70% identity over 100bp window.

The plant specific RS and RS2Z subfamilies associate with the R2R3-MYB class within the network built from *Arabidopsis* genes demonstrating conserved AS with *Amborella* and at least one species of monocots. Many R2R3-MYB genes are represented in this network and possess plant-specific functions (Table S6). Li et al. (2006) identified conserved AS events within MYB59 and MYB48. Both genes are represented within our network and associate with SR proteins via a proline-rich spliceosome-associated protein AT4G21660 (Figure 6). Together, these results suggest a conserved connection between splicing factors and R2R3-MYB transcription factors. These two protein classes may cooperate during developmental processes and stress response in plants.

### Grass Lineage Enriched for Conserved Alternative Splicing

Within monocots we identified lineage-specific conserved AS events. There are 204 conserved AS events between banana and African oil palm that are absent in the grass species examined, likely lost in the grass family, and 239 conserved AS events conserved in banana, African oil palm and at least one grass species. The number of conserved AS between two species in the PACMAD clade (maize, sorghum, and foxtail millet) is similar to the number of conserved AS events between two species within the BEP clade (rice and *Brachypodium*). 224 conserved AS events were found conserved across five grass species examined (maize, sorghum, foxtail millet, rice, and *Brachypodium*). In addition, the proportion of genes within syntenic blocks with AS events in the grasses is approximately twice that compared to those outside of syntenic blocks, and this trend is also mirrored across those genes with conserved AS events (Figure 3).

In general, the average values of Ka/Ks across different AS events are less than 1, which suggests that the flanking exonic regions are undergoing purifying selection for conserved AS events. In contrast, previous studies have suggested only moderate selection pressure on the alternative vs. the constitutive regions (Xing and Lee 2006, Chen et al. 2006, Xing and Lee 2005). However, the selection on the flanking region of conserved AS is less explored, particularly in plants. In intron retention, alternative acceptor and exon skipping, there is no significant difference in Ka/Ks between flanking upstream and downstream exonic regions (Figure 4); however, we did observe significantly higher Ka/Ks in the upstream compared to downstream flanking regions in alternative donor and alternative terminal exon. These results indicate that the flanking downstream regions have stronger purifying selection compared to the upstream regions in AltD and ATE events, which suggest that a proportion of flanking upstream and downstream exonic regions of conserved AS events might harbor splicing regulatory elements.

We observed both conserved and subfunctionalization patterns of AS within SR protein encoding genes after the whole genome duplication event in the maize lineage. The plant-specific RS subfamily in maize and sorghum harbor conserved exon skipping although the maize orthologue has been reduced to a single copy after whole genome duplication (Figure 5A). In the RS2Z family, Maize has retained both homeologous copies of the RS2Z subfamily genes. In one case, it has maintained the conserved AS events in both maize homeologues including exon skipping and intron retention events (Figure 5B), however, in another case, the AS events have diverged between the sorghum gene and its two maize homoelogous (Figure 5B).

### Important Functional Conserved AS in SR Protein in Response to Environment Stress

Many stress responsive TFs have conserved AS. We focused on identifying conserved AS in stress responsive TFs during five major stresses: ABA, salt (NaCl), drought, light, and cold. Two genes AT1G78070 (Transducin/WD40 repeat-like superfamily protein) and AT4G27410 (RD26) are expressed during all five stresses. RD26 encodes a NAC transcription factor induced in response to desiccation. In our data, RD26 has conserved AS in 7 out of the 9 species we examined (except sorghum and *Brachypodium*), however we don’t have data to test whether this conserved AS is stress responsive. As illustrated in Figure S4, the exonic flanking sequence of this intron retention event has the highest sequence conservation across species than any other region of the gene, although there remains some similarity between maize and sorghum in the conserved intron region. This conserved intron retention event is likely broadly conserved across angiosperms.

Within the protein-protein interaction network based on *Arabidopsis* stress response TFs with conserved AS events identified in our data sets, we detect phosphatases, kinases and several ubiquitin genes in the center of the network (Figure 6). This result is in line with the recent discovery that alternative exons and exitrons (exon-like introns) have more phosphorylation sites and ubiquitination sites compared to constitutive exons (Marquez et al. 2015). In these conserved AS genes across the angiosperms, AS may be involved in signaling pathways responsible for de/phosphorylation and protein degradation pathways, which suggests an important regulatory role in plants. Furthermore, we detected an association between plant-specific RS2Z and RS subfamily splicing proteins and R2R3-MYB proteins via proline-rich spliceosome-associated proteins, suggesting the R2R3-MYB family and splice regulators may interact during abiotic stress responses (Figure 6). Another interesting finding in the network is SPA3 (Figure 6). In moss, red and blue light photoreceptors were shown to regulate AS and intron retention was mis-regulated in moss mutants defective in the red light sensing phytochromes (Wu et al. 2014), which suggests an ancient connection between light regulation and conserved AS in land plants. We found both conserved alternative donor and intron retention events in SPA3, which acts as a negative regulator during light signaling via suppression of photomorphogenesis (Laubinger and Hoecker 2003). The truncated protein produced by two AS events in SPA3 will properly interact with COPI but loses the ability to bind to DDB1 (Shikata et al. 2014). SPA3 is connected to 9 additional kinase genes in the network, all of which have conserved AS across the angiosperms (Figure 6). This suggests that AS of kinases might play an important role in the regulation of the light signal, perhaps by affecting a kinase mediated signal cascade linked to SPA3. Inclusive, our study suggests the conserved AS landscape in plants is complicated and needs further functional study to link the phenotype to conserved AS events and identify the regulatory code for functional relevant AS.

## Authors’ Contributions

WM, WBB designed the work. WM, WBB, LB, GF, JCS analyzed the data. WM and WBB wrote the manuscript with input from LB, GF and JCS.

## Acknowledgements

We thank Daniel Gates and Emily Josephs provide helpful comments for the manuscript. This work was supported by Department of Biological Sciences at University of Florida, Florida Genetics Institute, Graduate Student Fellowship and College of Liberal Arts and Sciences Dissertation Fellowship from University of Florida awarded to WM and National Science Foundation grants IOS-0922742 & IOS-1547787 (WBB).

